# Loss of PIKfyve in Rod Photoreceptors and RPE Cells Leads to Endolysosomal Dysfunction and Retinal Degeneration

**DOI:** 10.1101/2025.09.15.676371

**Authors:** Ammaji Rajala, Larissa J. Trevino, Thamaraiselvi Saravanan, Tyler M. Black, Akbar M. Bhat, Tuan Ngo, Mark Eminhizer, Jianhai Du, Visvanathan Ramamurthy, Raju V.S. Rajala

## Abstract

Photoreceptor outer segment (OS) degradation is primarily mediated by retinal pigment epithelial (RPE) cells through daily phagocytosis of shed distal OS tips. In contrast, much less is understood about the cell-autonomous mechanisms photoreceptors use to clear mislocalized molecules caused by protein misfolding or trafficking defects. Mislocalized or excess rhodopsin that fails to reach the OS is retained in the inner segment or cell body, where it is presumably degraded via the endolysosomal system. We identify PIKfyve, a phosphoinositide kinase that generates PI(3,5)P₂, as a key regulator of this pathway. Using Translating Ribosome Affinity Purification (TRAP), we find that PIKfyve is highly expressed in rod photoreceptors. Rod-specific PIKfyve deletion causes progressive retinal degeneration, marked by inner segment vacuolation, elevated LAMP1/2, thinning of the outer nuclear layer, and eventual loss of rod and cone function. Loss of one copy of PIKfyve in rod photoreceptors accelerates degeneration in P23H rhodopsin mutant mice. In RPE cells, PIKfyve loss disrupts phagocytosis and autophagy, leading to accumulation of rhodopsin, LAMP1, LC3A/B, and lipid droplets, along with metabolic disturbances. These findings demonstrate that PIKfyve is essential for photoreceptor and RPE health by regulating lysosomal function, phagocytosis, autophagy and metabolism, and suggest that enhancing PIKfyve activity could be a therapeutic strategy for retinal degenerative diseases.

## Introduction

Phosphoinositides (PIPs) make up only a small part of cell phospholipids, but they influence nearly all aspects of a cell’s life and death[1–5]. Seven PIPs are produced through reversible phosphorylation in cells, regulating many processes including membrane budding and fusion, cytoskeletal assembly, vesicular transport, ciliogenesis, and signal transduction[1–6]. Mutations in phosphoinositide (PI)-converting enzymes lead to severe neurological and ophthalmological defects [7–11]. Human genetic diseases have been linked to genes that encode PI-converting enzymes and their effector proteins[11–16], which are implicated in neurological and ocular disorders, although the precise role of PIP metabolism imbalance in these diseases remains poorly understood.

In the mammalian genome, there are approximately 50 genes encoding phosphoinositide (PI) kinases and phosphatases, which generate seven distinct PIPs [17]. Their distribution in retinal cell types was previously unknown. Using translating ribosome affinity purification (TRAP), we mapped the expression of PI-converting enzymes across retinal cells, revealing enrichment of PI-converting enzymes in retinal neurons, and our data indicate that these enzymes are enriched in retinal neurons (rods, cones, RGCs) and depleted in Müller glia and RPE[18]. Our data also indicate that one of the PI kinases, PIKfyve, is significantly enriched in rod and cone photoreceptor cells compared to RGC, Müller glia, and RPE cells. PIKfyve, a phosphoinositide 5-kinase, catalyzes the phosphorylation of phosphatidylinositol 3-phosphate (PI(3)P) to produce phosphatidylinositol 3,5-bisphosphate (PI(3,5)P_2_) and PI to PI(5)P [19]. PIKfyve plays a key role in converting early endosomes into late endosomes and lysosomes, processes vital for breaking down cellular waste and recycling materials [20, 21]. Mutations or disruptions in PIKfyve activity or its associated proteins are linked to neurological disorders characterized by enlarged lysosomes and impaired lysosomal function [7, 22–26]. In humans, heterozygous mutations in *PIKFYVE* have been linked to fleck corneal dystrophy and congenital cataracts [10, 27]. The lipid PI(3)P is the substrate of PIKFyve, and loss of PI(3) P-generating enzyme vps34 in rods leads to retinal degeneration as a result of defects in endosome recycling [28]. Its loss in RPE results in an inability of lysosomes to fuse with phagosomes, suggesting a key role in membrane fusion with lysosomes [29]. The PIKfyve–FIG4–VAC14 complex activates PIKfyve by stabilizing the kinase and promoting PI(3,5)P₂ synthesis [30]. VAC14 acts as a scaffold, bringing PIKfyve and FIG4 together, while FIG4 supports PIKfyve activation through complex formation rather than its phosphatase activity [30].

The degradation of photoreceptor outer segment (OS) components is mainly dependent on RPE cells, which perform daily phagocytosis of shed distal OS tips [31, 32]. Currently, the retina field uses the term trogocytosis instead of phagocytosis since the RPE cells nibble a small portion of outer segments (10%) instead of engulfing the entire photoreceptor cells [30]. In *Drosophila,* mislocalized rhodopsin is targeted for degradation via the endolysosomal system [33, 34]. *Drosophila* do not have RPE, and defects in the endolysosomal pathway impair lysosomal function and contribute to retinal degeneration [33, 34]. In contrast, much less is understood about the cell-autonomous mechanisms photoreceptors use to clear mislocalized molecules, whether due to protein misfolding or trafficking defects. Mislocalized or excess rhodopsin that fails to reach the OS is retained in the inner segment or cell body, where it is presumably targeted for degradation via the endolysosomal system. It has been shown previously that visual arrestin (ARR1) helps internalize rhodopsin through clathrin adaptor protein, AP-2, into the endolysosomal pathway. [35]. In K296E mutant rhodopsin, abnormal ARR1–AP-2 complexes misroute rhodopsin and trigger photoreceptor cell death [35]. Our current study indicates that PIKfyve regulates the endolysosomal pathway in PR and RPE, and loss of this pathway in PR contributes to PR degeneration.

## Results

### Enrichment of the FYVE finger-containing phosphoinositide kinase (PIKfyve) in rod photoreceptor cells

Using translating ribosome affinity purification (TRAP), we investigated the expression patterns of PI-converting enzymes specifically in retinal cells (Fig. 1A). We found that FYVE finger-containing phosphoinositide kinase (PIKfyve) was significantly enriched in rod, cone and RPE cells compared to total retina, Müller glia, and retinal ganglion cells (Fig. 1B). PIKfyve is a PI-5-kinase that phosphorylates PI to PI(5)P and PI(3)P to PI(3,5)P_2_ (Fig. 1C). The lipid PI(3,5)P_2_ has previously been shown to be involved in the biogenesis (involves fusion/fission and resolution) of late endosomes and lysosomes [36], whereas PI(5)P has been suggested to have a role as a nuclear phosphoinositide in the regulation of gene expression [37]. To confirm the rod-specific expression of PIKfyve, we performed Opti-Prep density gradient centrifugation on an 8-40% gradient (Fig. 1D) to isolate rod outer segments (ROS) and broken inner segments, which release soluble cytoplasmic proteins [38]. The fractionation showed no significant co-migration between rhodopsin and PIKfyve, indicating that PIKfyve is localized in the inner segment, where most PI(3,5)P_2_ synthesis probably occurs (Fig. 1E). Our study demonstrates that PIKfyve is present in rod photoreceptor cells.

**Figure 1.**
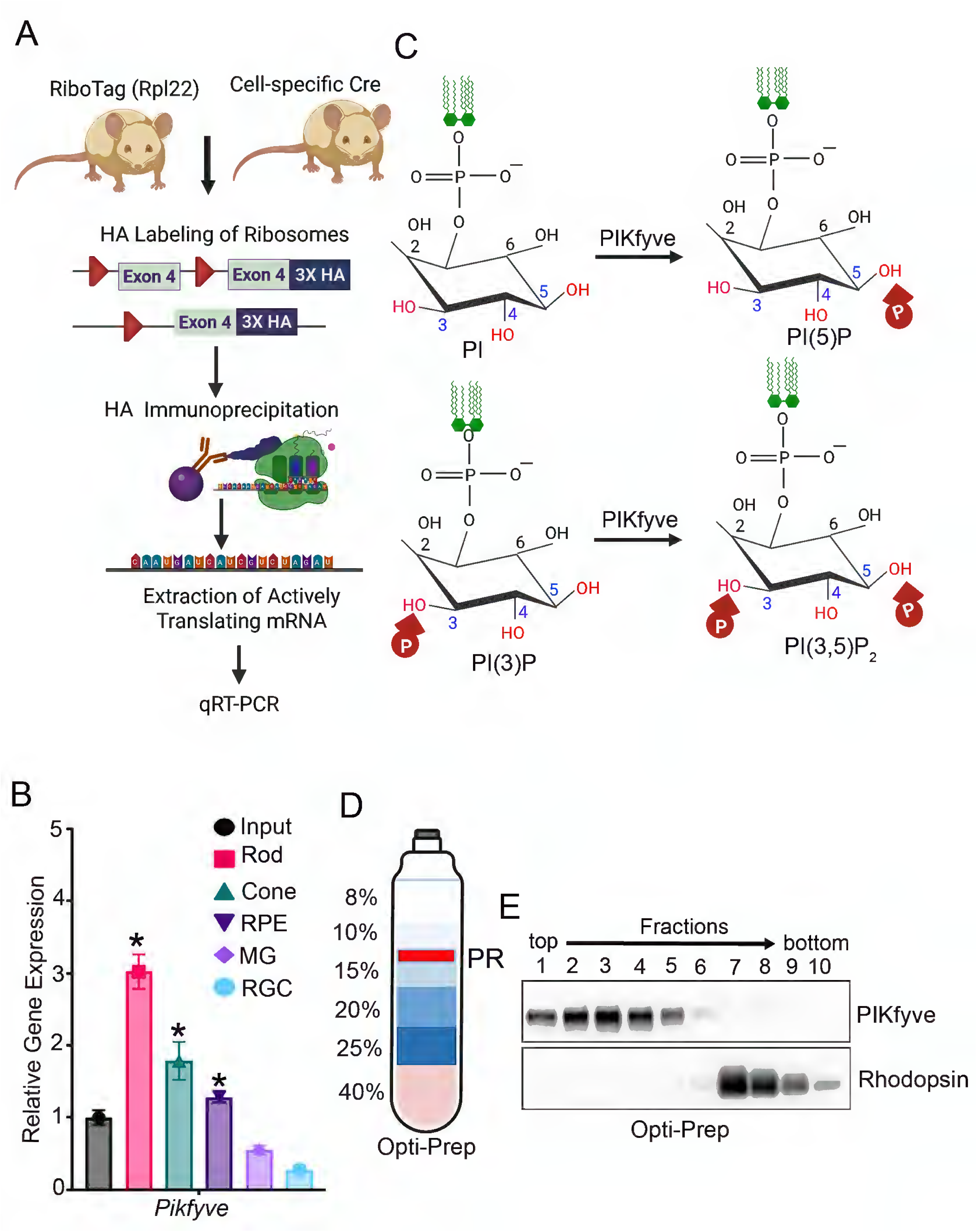
Expression of PIKfyve in retinal cells. A schematic diagram of Translating Ribosome Affinity Purification (TRAP). Rpl22 mice were bred with mice expressing Cre in rods, cones, RPE, Müller glia, and RGCs, followed by polyribosome immunoprecipitation. The resulting mRNA was isolated, and the expression of PIKfyve was measured using quantitative real-time PCR (qRT-PCR) (A). qRT-PCR analysis of mRNAs from rods, cones, RPE, Müller glia, and RGC using mouse *Pikfyve* primers, normalized to *Rpl37* and *Rpl38*. Data are presented as mean ± *SEM (n=3)*, with significance evaluated relative to Input levels (B). PIKfyve is a phosphoinositide 5-kinase that phosphorylates PI to PI(5)P and PI(3)P to PI(3, 5)P_2_ (C). An 8-40% Opti-Prep density gradient (D) centrifugation was used to isolate rod outer segments (ROS) and broken inner segments, which release soluble cytoplasmic proteins. Each fraction was analyzed by immunoblotting for PIKfyve and rhodopsin (E).

### Rod-photoreceptor-specific deletion of PIKfyve and its impact on rod structure and function

PIKfyve is essential for early embryonic development, and the global deletion of PIKfyve is embryonically lethal [39]. Thus, we have generated rod-specific PIKfyve knockout (*Pikfyve^rodko^*) mice using the rhodopsin promoter to drive Cre recombinase expression in rod cells (Fig. 2A). Four-week-old *Pikfyve^rodko^* mouse retina shows a significant reduction in PIKfyve protein compared to control (*Pikfyve^flox/flox^*) mouse retina (Fig. 2B). Recently, SnxA protein from the amoeba *Dictyostelium discoideum* has been identified as a highly selective PI(3,5)P_2_-binding protein and characterized as a reporter for PI(3,5)P_2_ in both *Dictyostelium* and mammalian cells [40]. We cloned and expressed SnxA as a fusion protein with an N-terminal maltose-binding protein (MBP) tag and a C-terminal rhodopsin 1D4 tag. The MBP tag enables purification of the fusion protein from bacteria by binding to amylose beads, while the ID4 tag is used to detect the interaction of the fusion protein with PI(3,5)P₂. A PIP array confirmed SnxA’s specific binding to PI(3,5)P_2_ (Fig. 2C). We used SnxA to measure PI(3,5)P_2_ and 2XHRS probes to assess PI(3)P levels [41] on ELISA in *Pikfyve^flox/flox^* (littermate control) and knockout mouse retinas. We found significantly increased PI(3)P and decreased PI(3,5)P_2_ levels in knockout mouse retinas compared to control mice (Fig. 2D). The PI(3,5)P_2_ levels decreased by 80%, while PI(3)P levels increased 2.5-fold compared to control (Fig. 2D). Histological analysis of four-week-old *Pikfyve^rodko^* mice showed no morphological changes compared to *Pikfyve^flox/flox^* mice (Fig. 2E, F). We assessed retinal function in four-week-old *Pikfyve^flox/flox^* and *Pikfyve^rodko^* mice using electroretinography (ERG). We found no significant differences in rod (scotopic a-wave and scotopic b-wave) or cone (photopic b-wave) function in *Pikfyve^rodko^*mice compared to *Pikfyve^flox/flox^* mice (Fig. 2G).

**Figure 2.**
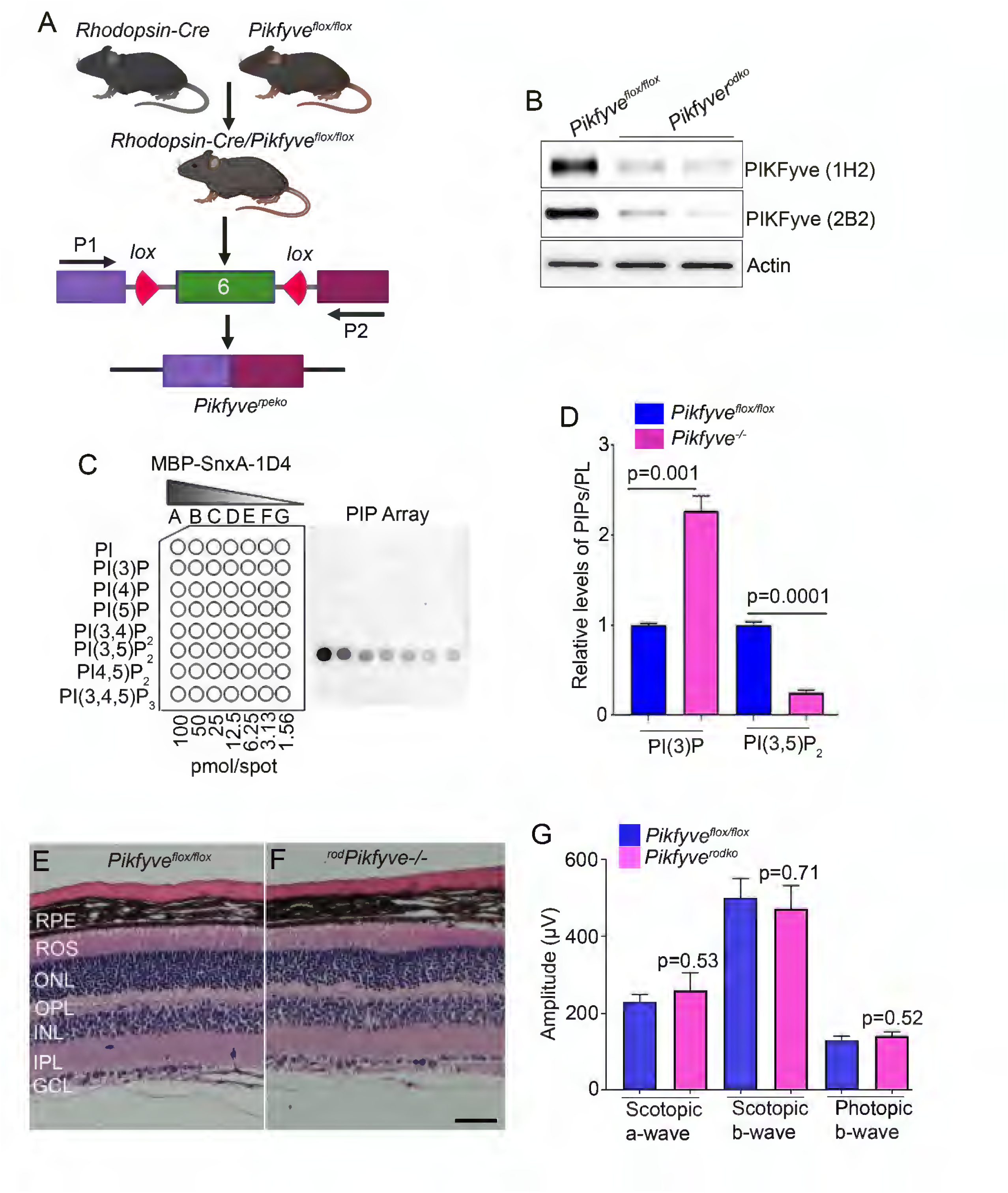
Generation of rod-specific PIKfyve KO mice and characterized biochemically, structurally, and functionally. Mice expressing floxed PIKfyve alleles flanking exon 6 were mated with mice expressing Cre-recombinase under the control of the rhodopsin promoter (A). P1 and P2 are primer sequences to identify desired genotypes. desired genotypes. Retina lysates from *Pikfyve^flox/flox^* and *Pikfyve^rodko^* mice were immunoblotted with PIKfyve (two different clones, IH2 and 2B2) and actin antibodies (B). PIP-Array demonstrates the specificity of the MBP-SnxA-1D4 probe to PI(3,5)P_2_ (C) and compares the levels of PI(3)P and PI(3,5)_2_ in control and PIKfyve KO mice (D). Four-week-old *Pikfyve^flox/flox^* and *Pikfyve^rodko^* mice retina sections ere stained with Hematoxylin and Eosin (H & E) (E, F). RPE, retinal pigment epithelium, ROS, rod outer segments, ONL, outer nuclear layer, OPL, outer plexiform layer, INL, inner nuclear layer, IPL, inner plexiform layer, GCL, ganglion cell layer. Electroretinography (ERG) was performed on 4-week-old *Pikfyve^flox/flox^* and *Pikfyve^rodko^* mice to examine rod (scotopic a- and scotopic b-wave amplitudes) and cone (photopic b-wave) function (G). Data are mean ± *SEM (n=12)*.

At 4 weeks, rod photoreceptor marker proteins, rhodopsin and rod arrestin levels remain unchanged in *Pikfyve^rodko^* retinas, which are comparable to *Pikfyve^flox/flox^* mice (Fig. 3A, B). However, we found significantly increased levels of lysosomal markers, LAMP1 and LAMP2, in *Pikfyve^rodko^* mouse retina compared to *Pikfyve^flox/flox^* mouse retina (Fig. 3C, D). To rule out whether the increased LAMP1 observed in immunoblots originates from Müller glia or microglia, retinal sections from four-week-old *Pikfyve^rodko^*and *Pikfyve^flox/flox^*mice were stained with a LAMP1 antibody. The results show increased LAMP1 expression in the external limiting membrane (ELM), located between the photoreceptor inner segments and Müller glial cells, in *Pikfyve^rodko^*mice compared to *Pikfyve^flox/flox^* mice (Fig. 3E, F). The punctate staining pattern of LAMP1 in the inner segment resembles the previously reported LAMP2 pattern in the same region [42].

**Figure 3.**
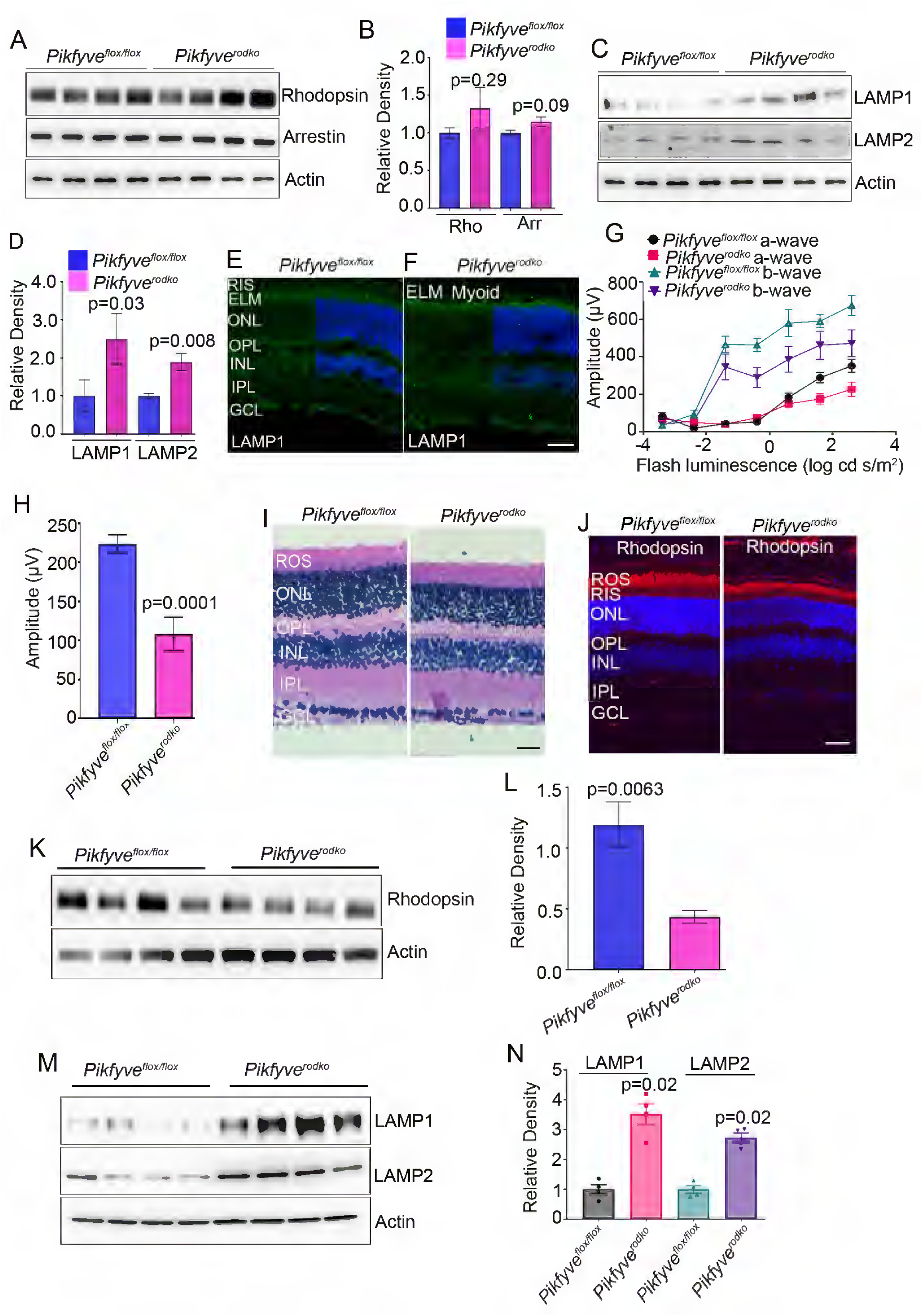
Age-dependent changes in rods lacking PIKfyve. Four-week-old *Pikfyve^flox/flox^* and *Pikfyve^rodko^* mouse retinal proteins were subjected to immunoblot analysis with rod photoreceptor markers (rhodopsin, arrestin) (**A**), lysosomal markers (LAMP1, LAMP2) (**C**), and normalized to actin (**B**, **D**). Data are mean ± *SEM (n=4).* Retinal sections from four-week-old *Pikfyve^flox/flox^* and *Pikfyve^rodko^* mice were stained with LAMP1 antibody (E, F). ERG was performed on 14-week-old *Pikfyve^flox/flox^* and *Pikfyve^rodko^* mice to examine rod (scotopic a- and scotopic b-wave amplitudes) (G) and cone (photopic b-wave) (H) function. Data are mean ± *SEM (n=8).* Morphological examination with H&E stain shows a decrease in ONL and a reduced ROS layer (I), decreased rhodopsin expression (J), and protein levels (K, L) in *Pikfyve^rodko^* mice compared to *Pikfyve^flox/flox^* mice. Sixteen-week-old *Pikfyve^flox/flox^* and *Pikfyv^rodko^* mouse retina lysates were immunoblotted with LAMP1, LAMP2, and actin antibodies (M). Densitometry analysis normalized to actin (N). Data are mean ± *SEM (n=4)*.

By 16 weeks, *Pikfyve^rodko^* mice exhibited significantly reduced scotopic a-wave and b-wave amplitudes (rod function) (Fig. 3G) and photopic b-wave amplitudes (cone function) compared to *Pikfyve^flox/flox^*mice (Fig. 3H). Additionally, they showed thinning of the outer nuclear layer (Fig. 3I), decreased rhodopsin expression (Fig. 3J), and lower rhodopsin levels (Fig. (Fig. 3K, L) compared to *Pikfyve^flox/flox^* mice. The LAMP1 and LAMP2 levels were significantly higher in 16-week-old *Pikfyve^rodko^*compared to *Pikfyve^flox/flox^* mouse retina (Fig. 3M, N), indicating lysosomal dysfunction.

### Deletion of PIKfyve in *Pikfyve^rodko^* results in vacuolation and lysosomal enlargement

To examine potential ultrastructural defects in *Pikfyve^rodko^* mice, we performed transmission electron microscopy (TEM) on retinal tissue from 14-week-old *Pikfyve^rodko^* and *Pikfyve^flox/flox^*mice. In *Pikfyve^flox/flox^* mouse retina, there were no anomalies in the outer segment and inner segment regions (Fig. 4A-C). Enlarged vacuoles were observed in the inner segments of rod photoreceptors in *Pikfyve^rodko^* retinas but were absent in *Pikfyve^flox/flox^* mice (Fig. 4E, F). At higher magnification, double-membraned structures—presumably autophagosomes and some structures resembling uncleared lysosomes—were clearly visible in *^rod^Pikfyve^-/-^*retinas (Fig. 4G, F).

**Figure 4.**
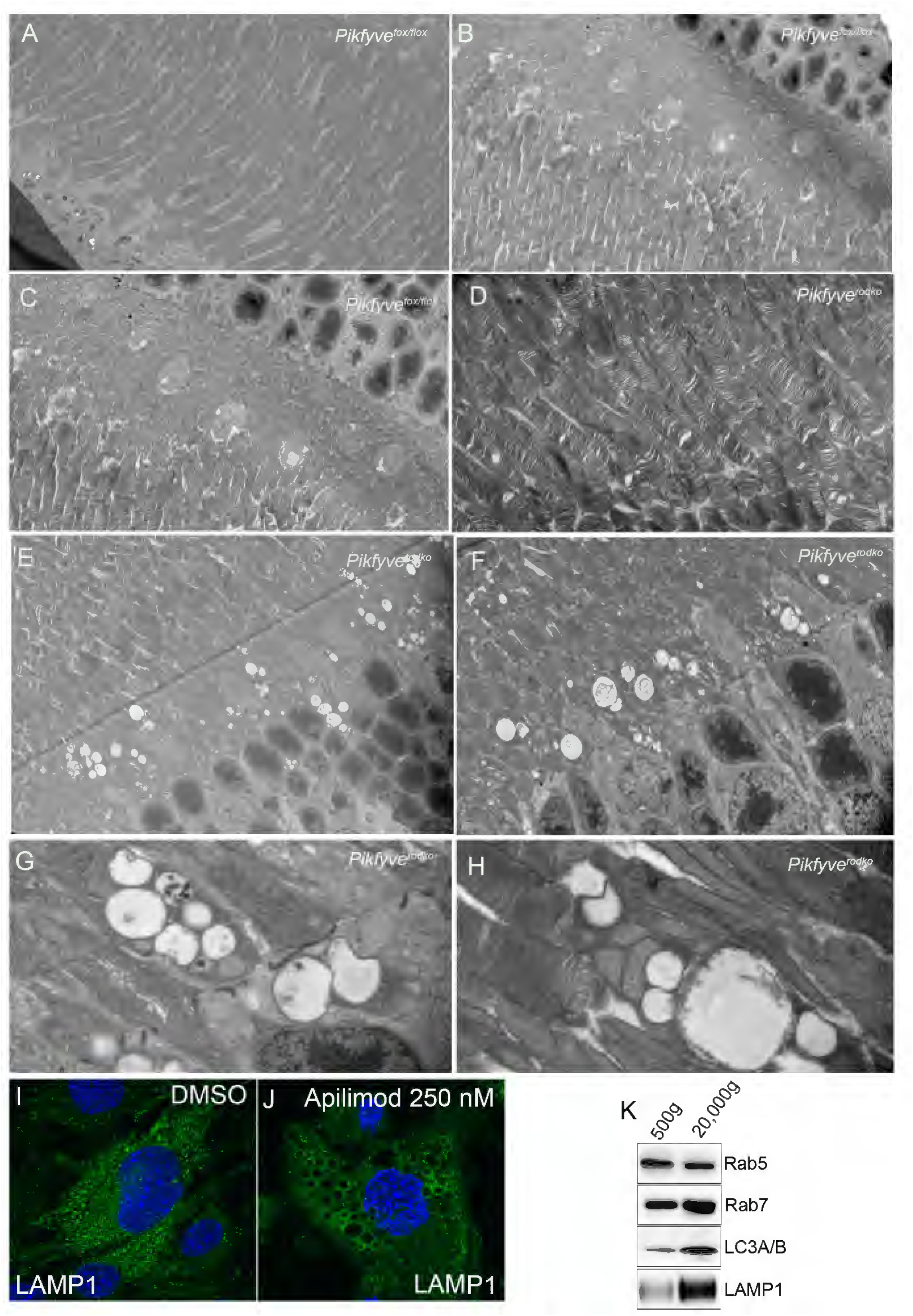
Abnormal vacuolization in *Pikfyve^rodko^* mice. Mouse retinal tissues from a 14-week-old *Pikfyve^flox/flox^* (A-C) and *Pikfyve^rodko^* (D-E) were analyzed by TEM. Panels F-H display a higher magnification of the region shown in Panel E. DMSO- and Apilimod-treated 661W cells were stained with LAMP1 antibody (I, J). Isolated vacuoles were immunoblotted using Rab5, Rab7, LC3A/B, and LAMP1 antibodies (K).

To investigate the contents of enlarged vacuoles caused by PIKfyve deletion, we treated 661W cells, derived from mouse cone photoreceptor cells [43], with the PIKfyve inhibitor Apilimod for 60 minutes at 37°C. After treatment, both DMSO control and Apilimod-treated cells were fixed and stained with a LAMP1 antibody. The results showed that the enlarged vacuoles in the treated cells were positively stained with LAMP1. (Fig. 4 I, J). LAMP1 is commonly recognized as a lysosomal marker, but it can also be found on late endosomes, phagosomes, and autophagosomes [30]. We isolated vacuoles from 661W cells treated with Apilimod using a method that involves sorbitol to loosen cell membranes [44]. Immunoblot analysis of the input fraction (500g) and isolated vacuoles (20,000g) was performed using markers for early endosomes (Rab5), late endosomes (Rab7), autophagy (LC3A/B), and lysosomes (LAMP1). The results showed enrichment of Rab7 and LC3A/B in the vacuoles compared to the input, indicating the presence of amphisomes (late endosomes and autophagosomes). Additionally, LAMP1 levels were significantly higher in the vacuoles than in the input (Fig. 4K). These findings suggest that the absence of PIKfyve impairs the fusion of late endosomes and autophagosomes with lysosomes.

### Loss of PIKfyve in rods exacerbates retinal degeneration in P23H rhodopsin mutant mice

P23H rhodopsin mutant mice are a widely used animal model for autosomal dominant retinitis pigmentosa (adRP) [45]. This model features a proline-to-histidine substitution at position 23 (P23H) in the rhodopsin (RHO) gene—a common mutation associated with autosomal dominant retinitis pigmentosa (adRP) in humans [45]. Morphological analysis of the retina revealed that at postnatal day 63 (P63), the ONL consisted of 9–10 rows of nuclei in WT, 6–7 rows in P23H heterozygous, and 0–1 row in homozygous P23H mice [45]. To assess whether reduced PIKfyve increases the susceptibility of already vulnerable photoreceptors to degeneration, and to investigate the role of PIKfyve in rod degeneration in P23H mice, we crossed P23H homozygous mice with *Pikfyve^rodko^* mice to generate *P23H^het^*and *P23H^het^*/*Pikfyve^rodhet^* mice (Fig. 5A). When we examined rhodopsin levels, we observed an absence of rhodopsin, significantly lower levels of cone arrestin, and no change in rod arrestin levels in *P23H^het^/Pikfyve^rodhet^*mice compared to *P23H^het^* mice (Fig. 5B, C). TRAP on two-month-old *Pikfyve^rodhet^*and *Pikfyve^rodko^* mouse retinas, followed by RNA sequencing, shows no change in the mRNA expression levels of rhodopsin, rod arrestin, and cone arrestin between the two genotypes (Fig. 5D). Consistent with decreased levels of rhodopsin on immunoblotting, immunohistochemistry further confirms reduced expression of rhodopsin in *P23H^het^/Pikfyve^rodhet^*compared to *P23H^het^* mice. The outer nuclear layer (ONL) contained 11 to 12 rows of photoreceptor nuclei, which is consistent with the normal range observed in rodents lacking retinal degeneration [46]. In the one-month-old *P23H^het^/Pikfyve^rodhet^*mouse retina, we observed approximately 5-6 rows of photoreceptor nuclei, indicating retinal degeneration. These observations suggest that the loss of one copy of PIKfyve in rods of P23H mice results in the exacerbation of both rod and cone degeneration.

**Figure 5.**
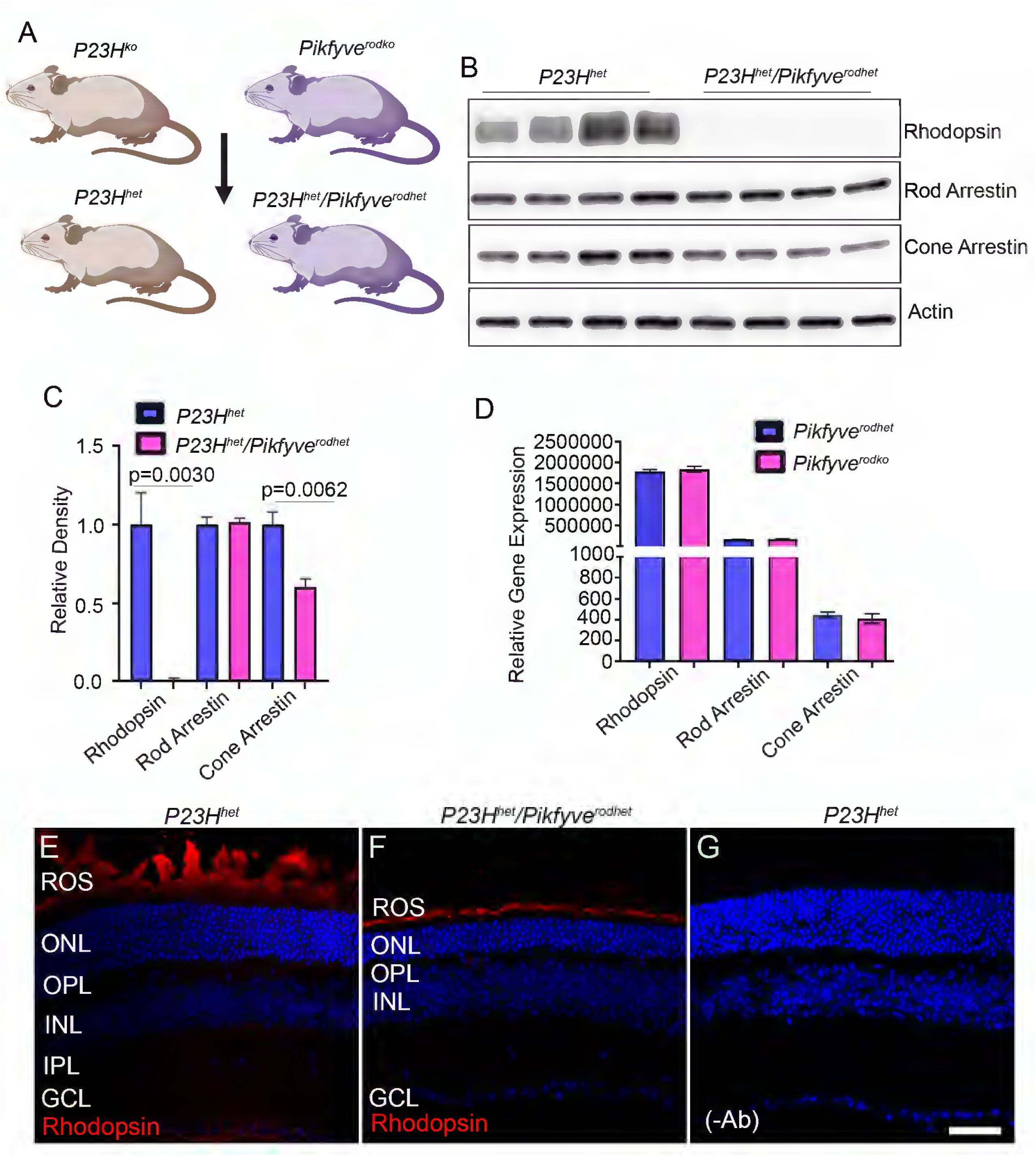
Loss of PIKfyve in rods exacerbates retinal degeneration in P23H rhodopsin mutant mice. We bred P23H homozygous mice with *Pikfyve^rodko^* mice to generate *P23H^het^* and *P23H^het^*/*Pikfyve^rodhet^* mice (A). Retina lysates from one-month-old *P23H^het^* mice and *P23H^het^/Pikfyve^rodhet^* mice were immunoblotted with rhodopsin, rod arrestin, cone arrestin, and actin antibodies (B), and their densities were normalized to actin (C). Data are mean ± *SEM*, (*n=4*). TRAP on two-month-old *Pikfyve^rodhet^* and *Pikfyve^rodko^* mouse retinas, followed by RNA sequencing, shows no change in the expression levels of rhodopsin, rod arrestin, and cone arrestin between the two genotypes (D). Data are mean ± *SEM*, (*n=3*). Retinal cryo sections from *P23H^het^* (E) and *P23H^het^/Pikfyve^rodhet^* (F) mice were stained with rhodopsin antibody. Panel G shows the omission of the primary antibody.

### Studies on PIKfyve deletion in the retinal pigment epithelium

The distal tips of the outer segments are phagocytosed daily by the retinal pigment epithelium (RPE). Our TRAP data also show the expression of *PIKfyve* in the RPE. Thus, we deleted PIKfyve in the RPE using Cre under the control of the RPE-specific bestrophin promoter (Fig. 6A) and examined the effect of PIKfyve loss on the RPE. Flat mounts were prepared from two-month-old RPE-specific PIKfyve KO mice (*Pikfyve^rpeko^*) and littermate controls (*Pikfyve^flox/flox^*) and stained with rhodopsin, LAMP1, and LC3A/B antibodies. The results indicate increased rhodopsin (Fig. 6B, C), LAMP1 (Fig. 6D, E), and LC3A/B (Fig. 6F, G) staining in *Pikfyve^rpeko^* mouse RPE compared to *Pikfyve^flox/flox^*mice. The increased rhodopsin expression suggests a slowdown of RPE phagocytosis, and the increased LAMP1 and LC3A/B expression indicate a lysosomal autophagy and LC3-mediated phagocytosis defect. Oil Red O is a fat-soluble dye used for staining neutral lipids and triglycerides in cells and tissues [47]. Increased Oil Red O-positive staining correlates with disease severity, especially in models mimicking dry age-related macular degeneration (AMD) [48, 49]. We observed increased Oil Red O staining in the RPE of two-month-old *Pikfyve^rpeko^*mice compared to *Pikfyve^flox/flox^* mice (Fig. 6H-N). Our studies demonstrate that deleting PIKfyve in the RPE leads to increased staining for rhodopsin, LAMP1, and LC3A/B, indicating impaired phagocytosis and autophagy. Additionally, higher Oil Red O staining suggests lipid buildup related to dry AMD-like pathology.

**Figure 6.**
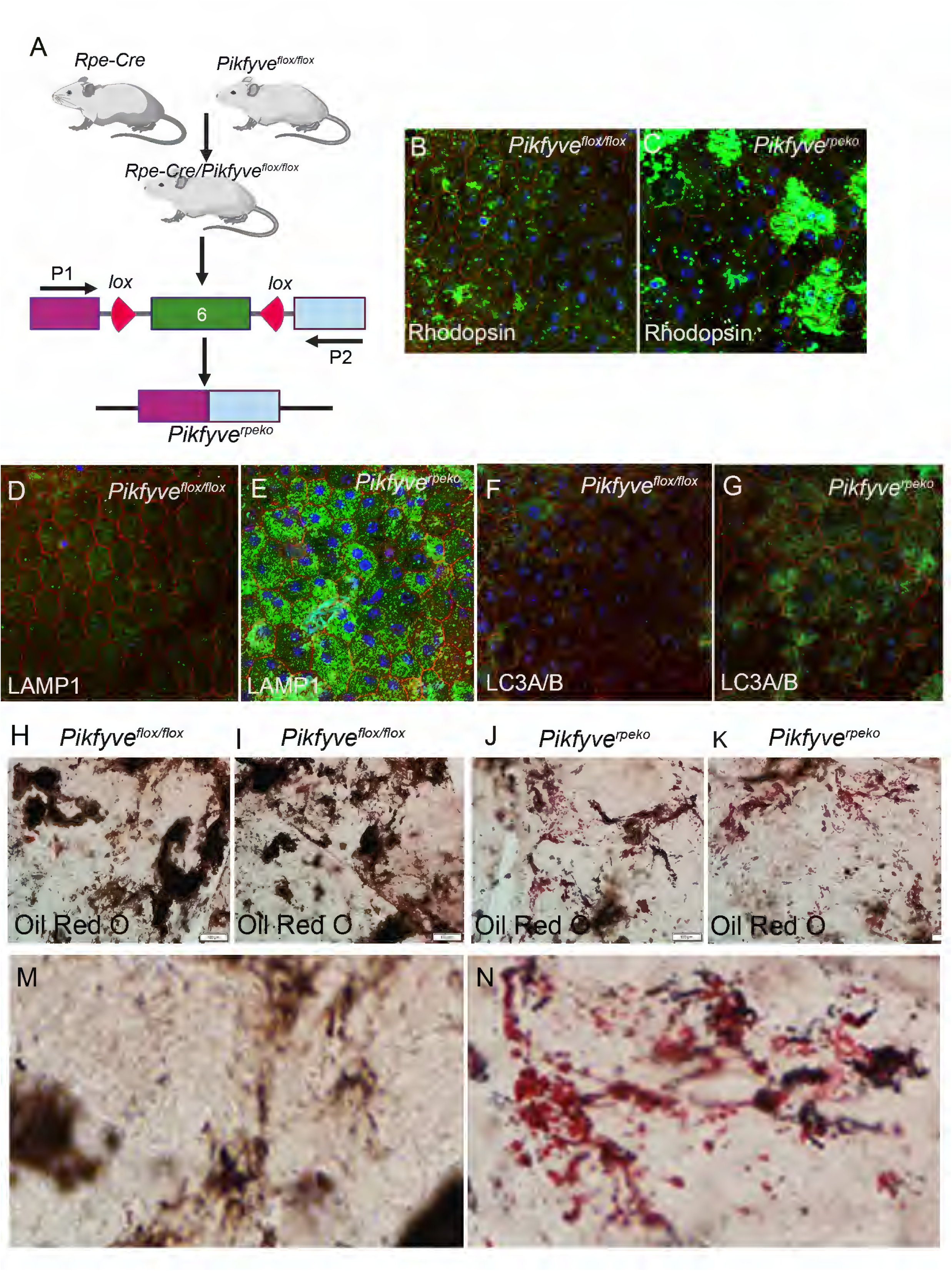
Generation and characterization of Rpe-specific PIKfyve knockout mice. Mice with floxed PIKfyve alleles flanking exon 6 were bred with mice expressing Cre recombinase under the control of the bestrophin (BEST1) promoter (A) to produce Rpe-specific PIKfyve KO mice (*Pikfyve^rpeko^*). P1 and P2 are primer sequences to identify desired genotypes. desired genotypes. Flat mounts were prepared from *Pikfyve^flox/flox^* and *Pikfyve^rpeko^* mice, then stained with antibodies for rhodopsin (B, C), LAMP1 (D, E), and LC3A/B (F, G). Oil Red O staining of retinal flatmounts from two-month-old *Pikfyve^flox/flox^* and *Pikfyve^rpeko^* mice (H-K). Panels M and N show higher magnifications of panels H (*Pikfyve^flox/flox^*) and K (*Pikfyve^rpeko^*).

### Effect of PIKfyve loss on retinal metabolism

Muscle-specific Pikfyve gene disruption has been shown to cause changes in energy metabolism [50]. To evaluate how PIKfyve deficiency in rods affects retinal metabolism, we analyzed steady-state metabolite levels in 8-week-old *Pikfyve^flox/flox^* and *Pikfyve^rodko^*mouse retinas using LC-MS (Fig. 7A). Principal component analysis (PCA) of the samples showed a clear separation between the *Pikfyve^flox/flox^* and *Pikfyve^rodko^* mouse retinas (Fig. 7B). Our study showed that several metabolites were altered in *Pikfyve^rodko^* mouse retinas compared to *Pikfyve^flox/flox^* mouse retinas (Fig. 7C).

**Figure 7.**
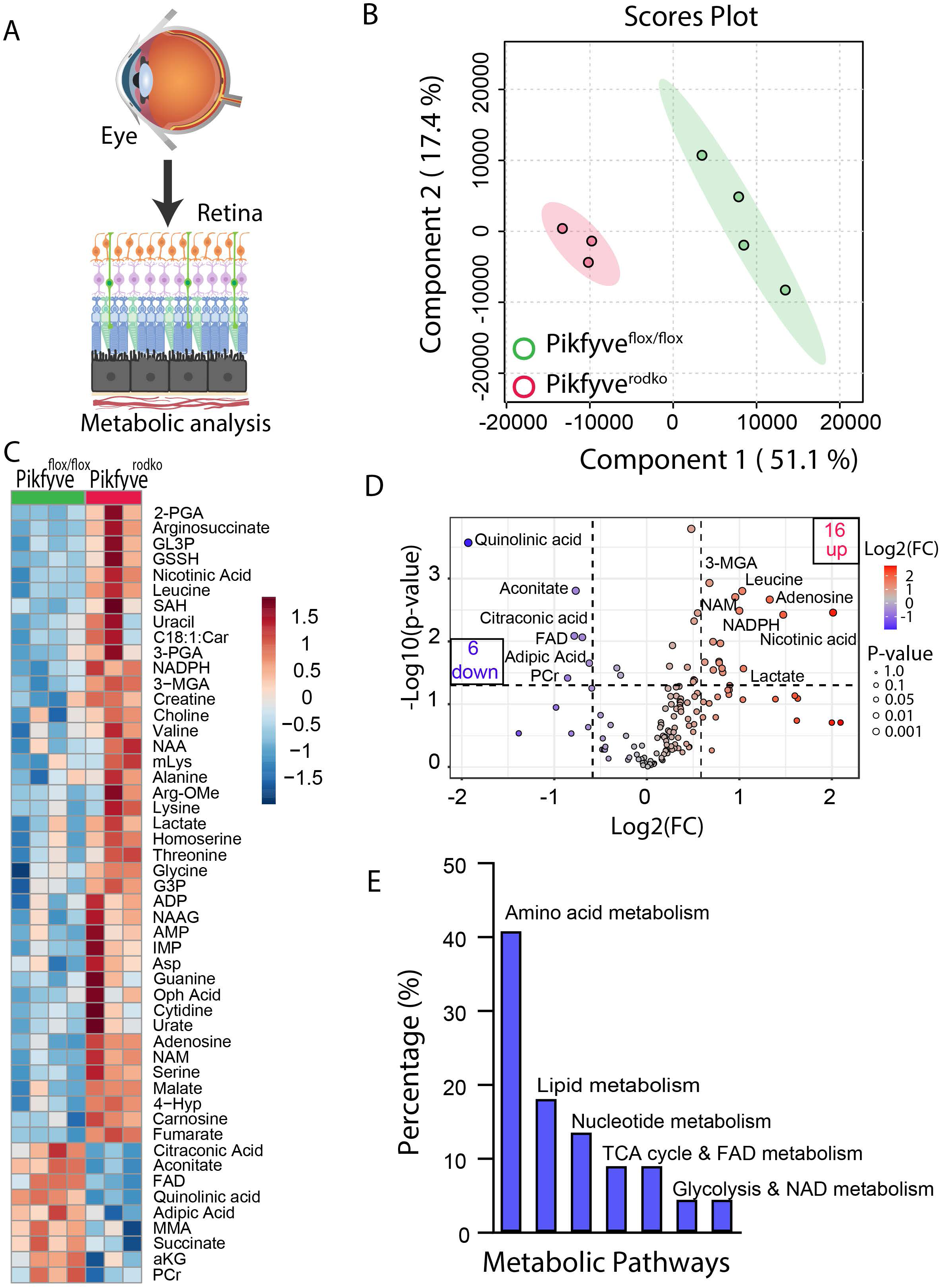
Impact of PIKfyve deficiency on retinal metabolism. Retinas were harvested from 8-week-old *Pikfyve^flox/flox^* and *Pikfyve^rodko^* mice to examine the steady-state levels of retinal metabolites (A). Principal component analysis (PCA) of the samples showed a clear separation between the *Pikfyve^flox/flox^* and *Pikfyve^rodko^* mouse retinas (B). A heatmap of changed metabolites in *Pikfyve^rodko^* compared to *Pikfyve^flox/flox^* mouse retina (C). Volcano plot showing differential metabolites in *Pikfyve^rodko^* compared to *Pikfyve^flox/flox^* mouse retina (D). Horizontal line indicates changes with (p < 0.05). Data are mean ± *SEM, (n = 4 WT; n=3 KO).* The histogram shows the pathway alterations in *Pikfyve^rodko^* compared to *Pikfyve^flox/flox^* mouse retina (E).

In the *Pikfyve^rodko^* mouse retina, we observed higher levels of certain metabolites compared to the *Pikfyve^flox/flox^*retina: 2-phosphoglyceric acid, L-argininosuccinic acid, glycerol 3-phosphate, oxidized glutathione, nicotinic acid, L-leucine, glyceraldehyde 3-phosphate, L-homocystine, NADPH, 3-methylglutaric acid, creatinine, adenosine, nicotinamide, L-serine, N-acetylaspartylglycine, malate, L-hydroxyproline, L-carnosine, and fumarate. Conversely, levels of citraconic acid, aconitate, FAD, quinolinic acid, succinate, and adipic acid were significantly reduced in *Pikfyve^rodko^* compared to *Pikfyve*^flox/flox^ mice. In contrast, phosphocreatine, succinate, and methylmalonate levels were decreased in *Pikfyve^rodko^* relative to *Pikfyve*^flox/flox^ mice (Fig. 7C, D). Based on the metabolomics data, the affected pathways in the retina where PIKFyve is deficient in rod photoreceptor cells include amino acid metabolism, lipid metabolism, nucleotide metabolism, the TCA cycle, and FAD metabolism, glycolysis, and NAD metabolism (Fig. 7E).

The retina is a highly metabolic tissue and depends exclusively on glycolysis, whereas RPE cells rely on oxidative phosphorylation [51]. Glucose is spared by RPE and is used by photoreceptor cells to produce lactate. Pyruvate kinase M2 is highly expressed in photoreceptor cells, while PKM1 is found in the inner retina [38]. PKM2 is allosterically activated by fructose 1,6-bisphosphate (FBP), which is cleaved by Aldolase to produce 3-carbon sugars, thereby regulating PKM2 activity [38]. On the other hand, PKM2 activity is inhibited by phosphorylation by oncogenic protein kinases [38]. LDHA is mainly expressed in photoreceptor cells, whereas LDHB is predominantly found in the RPE, where it catalyzes the conversion of lactate to pyruvate to fuel their mitochondria [52]. Glucose starvation has been shown to induce autophagy via ULK1-mediated activation of PIKfyve [53]. Thus, we examined the levels of these proteins in the RPE and retina of *Pikfyve^flox/flox^* and *Pikfyve^rpeko^*mice. We found significantly decreased levels of PKM2 and LDHA in two-month-old *Pikfyve^rpeko^* mouse RPE compared to *Pkm2 ^flox/flox^*mouse RPE (Fig. S1). The levels of PKM2 are also decreased in *Pikfyve^rpeko^*mouse RPE compared to *Pikfyve^flox/flox^* mouse; however, the difference is not statistically significant (Fig. S1A, D). We found significantly increased levels of ALDOC, Glut1, rod arrestin, and M-opsin in the retina of *Pikfyve^rpek^*° mice compared to *Pikfyve^flox/flo^*^x^ mice (Fig. S1B, C, E). The level of rhodopsin is also higher in the retina of *Pikfyve^rpeko^*mice compared to *Pikfyve2^flox/flo^*^x^ mice; however, the difference is not statistically significant. The increased rhodopsin in the retina could be due to decreased phagocytosis ability of RPE lacking Pikfyve.

## Discussion

This study first demonstrates that the loss of PIKfyve in rods leads to lysosomal dysfunction followed by retinal degeneration, while the loss of PIKfyve in the RPE results in the accumulation of lipid deposits. In *Drosophila*, defective rhodopsin trafficking leads to the accumulation of late endosomes and light-dependent degeneration, underscoring the importance of efficient endolysosomal degradation for photoreceptor survival [33, 34]. These findings also suggest that the endolysosomal degradative pathway is highly conserved in both invertebrates and vertebrates. Furthermore, the results highlight the critical role of PIKfyve in rod photoreceptors and the RPE.

In *Pikfyve^rodko^* mice, no mislocalization of rhodopsin was observed. Instead, our phenotype resembles that of a lysosomal storage disorder (LSDs), characterized by the formation of lysosomal storage vacuoles and elevated levels of lysosomal proteins [54, 55]. Increased plasma levels of LAMP-1 and LAMP-2 have been identified as diagnostic markers for individuals affected by LSDs [54, 55]. Our data suggest that the loss of PIKfyve in rod photoreceptors causes degeneration, likely due to lysosomal dysfunction, with the loss of PI(3,5)P_2_ disrupting lysosomal homeostasis, leading to enlargement and impaired function. These studies suggest that proteins in the inner segment, excess rhodopsin that is not targeted to the outer segment, or misfolded proteins, including rhodopsin [56], could be cleared through the endolysosomal pathway in rods. Consistent with this idea, losing one copy of PIKfyve in *P23H^het^* mice exacerbates photoreceptor degeneration by one month, emphasizing the role of PIKfyve in clearing misfolded proteins. Activation of PIKfyve could provide a therapeutic target for photoreceptor survival under disease conditions.

Previous studies have shown that vacuolization occurs in rods of P23H knock-in mice [57] and *Xenopus laevis* models expressing P23H rhodopsin, following light exposure [58]. The inositol-requiring enzyme 1 (IRE1) pathway facilitates the degradation of misfolded rhodopsin through proteasomal and lysosomal pathways without affecting wild-type rhodopsin [59]. The P23H mutation in rhodopsin causes protein misfolding and Endoplasmic Reticulum (ER) retention, leading to chronic ER stress, activation of the Unfolded Protein Response [59] and Endoplasmic Reticulum-Associated Degradation (ERAD), and eventual overload of cellular clearance pathways [57]. This results in rhodopsin aggregation, disrupted trafficking, oxidative and mitochondrial stress, and activation of multiple cell-death programs, culminating in progressive photoreceptor loss. A recent bioRxiv preprint shows that PIKfyve regulates ER-lysosome contacts to control ER morphology and motility. Inhibition of PIKfyve reduces ER reticulation and motility due to perinuclear lysosome clustering and hyper-tethering of the ER protein protrudin via excess PtdIns(3)P (Jenkins 2025).

Our ultrastructural studies reveal the accumulation of vacuoles in rod inner segments, and some of the structures appear to be double-membranous, suggestive of autophagosomes. Some of the structures could be uncleared lysosomes. Our data from vacuoles isolated from PIKfyve-inhibitor-treated 661W cells show increased Rab7 and LC3A/B levels, indicating an accumulation of amphisomes—hybrid organelles formed by the fusion of autophagosomes with late endosomes [60]. Amphisomes subsequently fuse with lysosomes to form autolysosomes or amphisomal lysosomes, facilitating degradation. Supporting this, *pikfyve* zebrafish mutants exhibit aberrant vacuolation, characterized by the accumulation of Rab7+LC3+ amphisomes in lens cells [10].

Apilimod is being studied for a wide range of diseases, including autoimmune disorders like Crohn’s disease [61] and psoriasis [62], neurodegenerative conditions such as ALS [63], various cancers [64], and viral infections like COVID-19 [65] and Ebola [66]. Its potential uses also include cardiovascular diseases. The drug mainly works by inhibiting PIKfyve activity involved in endolysosomal trafficking and cytokine signaling pathways. This inhibition can decrease inflammatory cytokine release and disrupt intracellular trafficking—key mechanisms for its antiviral and anticancer effects. However, despite these promising therapeutic prospects, Apilimod’s usage raises safety concerns. Blocking PIKfyve interferes with lysosomal function, leading to cellular vacuolization and lysosomal swelling, which impair protein degradation and autophagy. Immune suppression might also occur due to decreased T cell activity and lower cytokine production, possibly increasing the risk of infections. Especially vulnerable are RPE cells, which rely on intact endo-lysosomal pathways for phagocytosing photoreceptor outer segments; their dysfunction could contribute to retinal degeneration and vision loss.

RPE plays a vital role in preserving retinal health and vision. It aids photoreceptors by removing their shed outer segments, regenerates visual pigments via the visual cycle, and transports nutrients, ions, and waste between the retina and choroid. Additionally, the RPE constitutes part of the outer blood-retinal barrier and secretes key growth factors, such as VEGF and PEDF, to support retinal structure and function. These functions collectively ensure the survival of photoreceptors and proper visual function. The endo-lysosomal pathway plays a vital role in supporting the function and survival of RPE cells. This pathway is responsible for processing endosomes, phagosomes, and autophagosomes, all of which are essential for cellular clearance and homeostasis [67, 68]. Defective lysosomal function, as observed in lysosomal storage disorders, has been linked to progressive retinal degeneration [69]. In the RPE, loss of PIKfyve causes increased staining of rhodopsin, LAMP1, and LC3, indicating impaired phagocytosis, lysosomal activity, and autophagy. The elevated LAMP1 levels may indicate a defect in the fusion of lysosomes with endosomes, phagosomes, or autophagosomes. Although LAMP1 is primarily a lysosomal marker, it can also accumulate on late endosomes, phagosomes, and autophagosomes [70].

Earlier studies have demonstrated that oil red O binds lipids accumulated in Bruch’s membrane (BrM) in AMD patients [48]. Aging is the primary risk factor for AMD, and soft drusen, along with basal laminar deposits, are lipid-rich extracellular lesions specific to AMD. Oil Red O binding to neutral lipids indicates a significant age-related deposit in the BrM and is the first identified drusen component [49]. The RPE lacking PIKfyve shows increased Oil Red O staining, indicating the importance of PIKfyve in lipid clearance and its role in AMD prevention. Interestingly, PIKFYVE has previously been identified as one of the genes associated with AMD. [71]; however, further studies have not been pursued. Additional research is needed to examine age-related changes in PIKfyve and lysosomal dysfunction.

In the *Pikfyve^rodko^* mouse retina, metabolomic profiling revealed widespread changes across several key metabolic pathways, indicating significant cellular stress and dysfunction. Elevated levels of glycolytic intermediates such as 2-phosphoglyceric acid, glyceraldehyde 3-phosphate, and lactate, along with decreased succinate and phosphocreatine, suggest a shift toward glycolysis and impaired mitochondrial oxidative phosphorylation. Accumulation of TCA cycle intermediates like malate and fumarate, coupled with reduced aconitate and succinate, further supports mitochondrial dysfunction and disrupted energy production. Amino acid metabolism was also affected, as shown by increased levels of L-leucine, L-serine, L-argininosuccinic acid, and L-homocystine, reflecting altered protein turnover and nitrogen balance. Elevated levels of oxidized glutathione, glutathione, and NADPH indicate increased oxidative stress and an active antioxidant response. Additionally, changes in purine metabolism, such as elevated adenosine and nicotinamide, and decreased quinolinic acid and FAD, suggest a disruption of the NAD+/FAD cofactor balance and redox signaling. Variations in lipid-related metabolites, such as glycerol 3-phosphate, L-acetylcarnitine, and choline, indicate impaired membrane metabolism and lipid turnover (Fig. 7D). Overall, these metabolic disturbances suggest that PIKFyve deficiency leads to impaired mitochondrial function, oxidative stress, energy imbalance, and metabolic inflexibility, likely contributing to the accelerated photoreceptor degeneration observed in these mice.

Our findings highlight the crucial role of PIKfyve in regulating retinal energy metabolism. Metabolomic profiling revealed significant alterations in steady-state metabolite levels in Pikfyve-deficient retinas, including disruptions in purine metabolism —a pathway crucial for nucleotide synthesis and energy transfer. These alterations indicate broader metabolic stress and impaired nucleotide homeostasis in the absence of PIKfyve. Since the retina depends on aerobic glycolysis, the reduced levels of PKM2 and LDHA in the RPE of PIKfyve-deficient mice suggest compromised glycolytic activity and altered lactate production, which could disrupt the metabolic connection between photoreceptors and the RPE. Additionally, increased expression of Glut1 and ALDOC, along with higher levels of photoreceptor-specific proteins, may represent compensatory efforts to meet energy needs. Overall, these results emphasize the vital role of PIKfyve in maintaining metabolic balance in the retina, particularly through its regulation of glycolysis, purine metabolism, and nutrient-responsive signaling pathways. These results align with previous studies that have linked PIKfyve to nutrient sensing and autophagy, underscoring its broader function in supporting cellular energy demands through the regulation of glucose metabolism.

In summary, PIKfyve-produced PI(3,5)P₂ is vital for controlling the endolysosomal pathway in both photoreceptor and RPE cells. Our results suggest that activating PIKfyve may offer neuroprotection in retinal diseases.

## Materials and Methods

### Animals

All animal procedures complied with the ethical standards outlined in the ARVO Statement for the Use of Animals in Ophthalmic and Vision Research and adhered to the NIH Guide for the Care and Use of Laboratory Animals. Experimental protocols received approval from the Institutional Animal Care and Use Committee (IACUC) at the University of Oklahoma Health Sciences Center. Breeding colonies of BEST1-Cre (Jax #017557), RiboTag (Jax #029977), P23H (Jax #017628), and floxed Pikfyve (Jax #029331) mice were obtained from The Jackson Laboratory (Bar Harbor, ME). Rhodopsin-Cre mice were kindly provided by Dr. Ching-Kang Jason Chen at Baylor College of Medicine (Houston, TX). All mice were housed in our institutional animal facility under a controlled 12-hour light/dark cycle (40–60 lux). Animals were screened and confirmed negative for rd1 and rd8 retinal degeneration mutations. To reduce experimental bias, mice were randomly assigned to groups matched for sex, age, and genetic background. Litters were pooled to prevent litter-specific confounding effects.

### Translational Ribosome Affinity Purification

The generation of rod, cone, RPE, Müller glia, and RGC-specific/Rpl22 mice has been previously described. Polyribosomes engaged in active translation were isolated using a modified protocol based on Cleuren et al. [72]. Retinas were dissected (aged 2–4 months) and immediately transferred into DMEM supplemented with cycloheximide (100 µg/mL) to halt translation elongation. The tissues were incubated in this solution for 10 minutes, then rapidly flash-frozen in liquid nitrogen. Frozen retinas were mechanically ground using a hand-held homogenizer. The resulting tissue powder was resuspended in 200 µL of ice-cold polysome extraction buffer, composed of 50 mM Tris-HCl (pH 7.5), 100 mM KCl, 12 mM MgCl₂, 1% Igepal CA-630, 1 mM dithiothreitol (DTT), 200 U/mL RNaseOUT, 1 mg/mL heparin sodium salt, 100 µg/mL cycloheximide, and a protease inhibitor cocktail (EDTA-free) prepared in DEPC-treated water. The homogenate was gently mixed by pipetting and centrifuged at 15,000 RPM for 10 minutes at 4 °C to clarify the lysate. The supernatant was then incubated with 5 µL of purified mouse monoclonal HA antibody per 200 µL of lysate for 1 hour at 4 °C to allow binding of tagged polyribosomes. Subsequently, pre-equilibrated magnetic protein G beads were added and allowed to bind the antibody complexes during a 30-minute incubation at 4 °C. To remove nonspecific interactions, the beads were washed three times with high-salt wash buffer containing 50 mM Tris-HCl (pH 7.5), 300 mM KCl, 12 mM MgCl₂, 1% Igepal CA-630, 1 mM DTT, 200 U/mL RNaseOUT, 1 mg/mL heparin, 100 µg/mL cycloheximide, and EDTA-free protease inhibitors. RNA was then extracted directly from the immunoprecipitated complexes using 500 µL of TRIzol reagent, followed by purification with the PureLink RNA Mini Kit (Ambion, Carlsbad, CA). Complementary DNA (cDNA) synthesis was performed using the SuperScript III First-Strand Synthesis System (Invitrogen) according to the manufacturer’s protocol. The isolated cDNA was used to quantify gene expression using qRT-PCR. To quantify mouse *Pikfyve* transcript levels in various retinal cell types, qRT-PCR was performed using the following primers: sense, GATTCATCCGGATTCCTCAA, and antisense, TAGCCTGGGGACTGACAGAT. Transcript levels were normalized to two reference genes, *Rpl37* and *Rpl38*, with the primers: Rpl37 sense – CGGGACTGGTCGGATGAG and antisense – TCACGGAATCCATGTCTGAATC; Rpl38 sense – CGCCATGCCTCGGAAA and antisense – CCGCCGGGCTGTCAG. For RNA sequencing, RNA will be sent to the RNA sequencing facility at MedGenome Inc. (Foster City, CA).

### Generation of Rod- and RPE-Specific Pikfyve Knockout Mice

Rod- and RPE-specific conditional Pikfyve knockout mice were created using the Cre-loxP system. Mice with floxed Pikfyve alleles were crossed with transgenic mice expressing Cre recombinase under the control of the rod-specific rhodopsin promoter (to produce rod-Cre Pikfyve-KO) or the RPE-specific bestrophin promoter (to produce RPE-Cre Pikfyve-KO). In both models, Cre recombinase mediates the deletion of exon 6 of the Pikfyve gene, resulting in tissue-specific gene knockout. Genotyping was performed by PCR using DNA extracted from tail biopsies. Rod-Cre (opsin-Cre) was identified with primers: sense 5′-TCAGTGCCTGGAGTTGCGCTGTGG-3′ and antisense 5′-CTTAAAGGCCAGGGCCTGCTTGGC-3′, which amplify a 500 bp product. RPE-Cre was detected with a different primer set: sense [GGC ACT GGC CAC AGA GTC] and antisense [GGT GTA CGG TCA GTA AAT TGG A], amplifying a 158 bp product. Floxed Pikfyve alleles were identified using sense 5′-GCC TGA GTT CTG AGA GTG AGT G-3′ and antisense 5′-CAC TGG CTA TCT GGC ATC C-3′ primers, which produced a 421 bp product for the wild-type, both a 421 bp and a 500 bp product for heterozygotes, and a 500 bp product for the mutant allele.

### Transmission Electron Microscopy

For transmission electron microscopy (TEM), mouse eyes were immersion-fixed using a detailed, multi-day protocol. On Day 1, mice were sacrificed by brief CO₂ exposure followed by cervical dislocation. Eyes were enucleated using toothed forceps and curved scissors, rinsed in Milli-Q water to remove blood and hair, and transferred to a fixative solution. A 20G needle punctured the cornea, which was then incised to allow fixative penetration. Eyes were incubated in glass vials containing fixative at room temperature for 30–60 minutes without agitation. Afterwards, eyes were placed in a drop of sucrose buffer, and the cornea, iris, and lens were carefully removed to make eyecups. Eyecups were returned to fresh fixative and gently rolled at room temperature for two days. On Day 3, eyecups were either transferred into sucrose buffer for same-day shipping or stored in fixative at 4 ° C for later processing. For dissection, eyecups were cut into quadrants with the retina side up and trimmed into smaller trapezoid pieces. Samples were washed in 0. 0.1M cacodylate buffer, post-fixed in 2% osmium tetroxide for 1 hour on ice, washed again, then incubated overnight at 4 ° C in 1% uranyl acetate. On Day 4, samples were washed, dehydrated in graded ethanol on ice, treated with propylene oxide, and infiltrated with resin mixtures in increasing concentrations. Samples were embedded in silicone molds with 100% resin and cured at 60 ° C for several days. On Day 7 or later, cured resin blocks were trimmed into trapezoids with a glass knife under a dissection microscope and prepared for sectioning. Semithin sections (0. 0.5–2 µm) were cut with a diamond knife, floated on water in the knife boat, collected onto slides, stained with Richardson’ s stain, and examined under a microscope. On Day 9, ultrathin sections (50–100 nm) were cut using a diamond knife at ultra-thin settings. Sections were floated as ribbons on water, flattened using chloroform vapor, and carefully collected onto glow-discharged, formvar-coated grids. Grids were air-dried and inspected under a microscope for section integrity before imaging.

### Statistical Analysis

Sample sizes were calculated using power analyses. All statistical analyses were performed with GraphPad Prism version 7.3. No data points were excluded from any analysis. The selection of statistical tests was based on experimental design. Before analysis, the normality of each dataset was checked using multiple tests, including the Anderson-Darling, D’Agostino-Pearson, Shapiro-Wilk, and Kolmogorov-Smirnov tests, to determine if the data followed a normal distribution. For datasets that failed normality tests, non-parametric methods—specifically, multiple Mann-Whitney U tests—were used to compare the groups. To adjust for multiple comparisons and control the false discovery rate, the two-stage Benjamini-Krieger-Yekutieli procedure was applied with a Q value threshold of 1%. For normally distributed data, comparisons between two groups used parametric tests with Welch’s correction to account for unequal variances. Statistical significance was based on p-values. One-way ANOVA was used to compare three or more independent groups, while two-way ANOVA evaluated the interaction between two categorical variables on a continuous outcome.

### Other methods

Electroretinography (ERG), tissue histology, Western blotting, and immunohistochemical staining were performed following previously established protocols [52]. The antibodies used in this study are listed in Supplemental Table 1. Targeted metabolomics was carried out by LC-MS as described [73]. We added Nicotinamide-D4 as the internal standard for quality control.

## Acknowledgements

This study was supported by grants from the National Institutes of Health EY035282, EY031324 (to J. D.), EY032462 (to J. D.), Retinal Research Foundation (To J.D.), NEI Core grant EY021725, Presbyterian Health Foundation, and an unrestricted grant from Research to Prevent Blindness, Inc., awarded to the OUHSC Department of Ophthalmology. The authors gratefully acknowledge Dr. Jennifer Hocking (University of Alberta) for her insightful comments and critical reading of the manuscript.

## Author Contributions

RVSR conceived and designed the study. RVSR, AR, LT, and TB conducted experiments. Data analysis was performed by RVSR and AR. RVSR and AR interpreted the results. TS performed the TEM, and VR analyzed the TEM images. TN, ME, and JD analyzed the retinal metabolites and interpreted the data. RVSR drafted and wrote the manuscript, incorporating feedback and critical input from all co-authors. All authors reviewed and approved the final version of the manuscript.

---

**Figure S1. Expression of RPE and Photoreceptor Markers in Mice Lacking PIKfyve in the RPE.** Protein lysates from RPE (A) and retina (B, C) from *Pikfyve^flox/flox^* and *Pikfyve^rpeko^* were immunoblotted with ALDOC, PKM2, p=PKM2, PKM1, LDHA, LDHB, Rpe65, GS, Glut1, Rhodopsin, rod arrestin, GFAP, GS, M-opsin, cone arrestin, and antibodies. Protein densities from RPE (D) and retina (E) were normalized to actin. Data are mean ± *SEM, (n = 4)*.

