## Supplementary figures and images for "Loss of PIKfyve in Rod Photoreceptors and RPE Cells Leads to Endolysosomal Dysfunction and Retinal Degeneration"

### Supplemental Figure 1

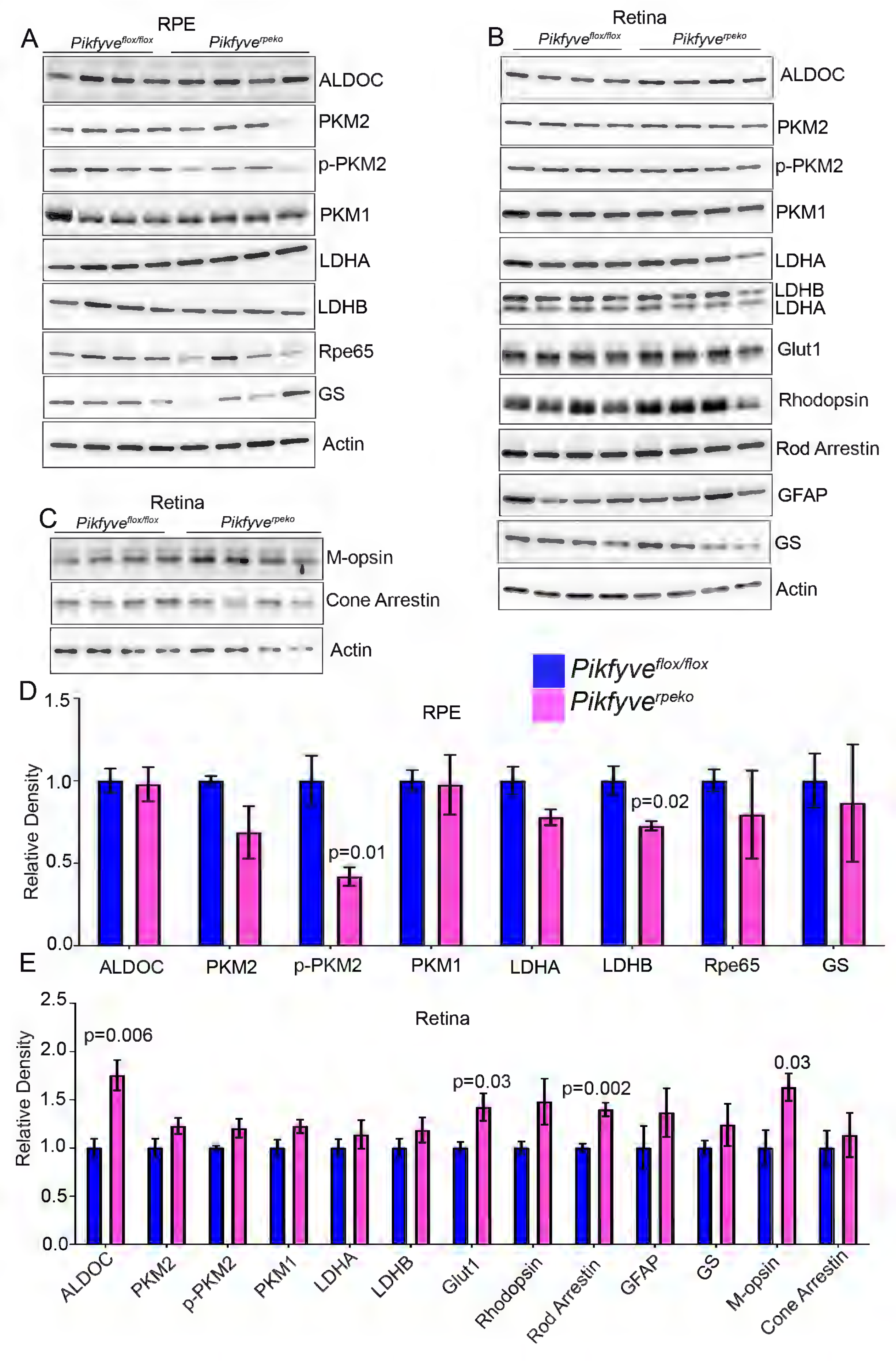
