## Supplemental Table 1 for "Loss of PIKfyve in Rod Photoreceptors and RPE Cells Leads to Endolysosomal Dysfunction and Retinal Degeneration"

| Table S1. Antibodies used for immunoblotting and immunofluorescence (IF) Analysis | | | | |
|---|---|---|---|---|
| **Antibody** | **Host species** | **Dilution** | **Manufacturer** | **Catalog number** |
| PIKFyve | Mouse | 1:1000 | Developmental Studies Hybridoma Bank (DSHB) | 1H2-s |
| PIKFyve | Mouse | 1:1000 | Developmental Studies Hybridoma Bank (DSHB) | 2B2-s |
| LDHA | Rabbit | 1:1000<br>1:50 (IF) | Proteintech | 21799-1-AP |
| LDHB | Rabbit | 1:1000<br>1:50 (IF) | Proteintech | 14824-1-AP |
| Pde6β | Mouse | 1:1000<br>1:25 (IF) | Santa Cruz | SC-377486 |
| Rhodopsin | Mouse | 1:1000<br>1:50 (IF) | In-house | Gift from Dr. Jim McGinnis (OUHSC) |
| Rod-Arrestin | Mouse | 1:1000<br>1:500 (IF) | In-house | Gift from Dr. Paul Hargrave (University of Florida) |
| M-opsin | Rabbit | 1:1000<br>1:100 (IF) | Millipore Sigma | AB5405 |
| S-opsin | Rabbit | 1: 100 (IF) | Millipore Sigma | ABN1660-1 |
| Cone-Arrestin | Rabbit | 1:1000 | Millipore Sigma | AB15282 |
| Actin | Mouse | 1:1000 (IB) | Thermo Fisher Scientific | MA1-744 |
| Glutamine synthetase (GS) | Mouse | 1:1000<br>1:50 (IF) | Abcam | Ab64613 |
| Glial fibrillary acid protein (GFAP) | Rabbit | 1:100 (IF) | Dako | 20334 |
| PKM1 | Rabbit mAb | 1:1000<br>1:50 (IF) | Cell Signaling Technology | 7067 |
| PKM2 | Rabbit mAb | 1:1000<br>1:100 (IF) | Cell Signaling Technology | 4053 |
| Phospho-PKM2 (Tyr105) | Rabbit | 1:1000<br>1:50 (IF) | Cell Signaling Technology | 3827 |
| Aldolase C | Mouse | 1:1000<br>1:100 (IF) | EnCor Biotechnology, Inc. | MCA-4A9 |
| Glut1 | Rabbit | 1:1000<br>1:25 (IF) | Novus Biologicals | NB110-39113 |
| Rpe65 | Rabbit | 1:1000 | In-house | Gift from Dr. Jian-Xing Ma |

| | | | | |
|---|---|---|---|---|
| Rab5A | Rabbit | 1:1000 | Cell Signaling Technology | 46449 |
| Rab7 | Rabbit | 1:1000 | Cell Signaling Technology | 9367 |
| LC3A/B | Rabbit | 1:1000 | Cell Signaling Technology | 12741 |
| LAMP1 | Rat | 1:1000<br>1:100 (IF) | Developmental Studies Hybridoma Bank (DSHB) | 1D4B |
| LAMP2 | Rat | 1:1000<br>1:100 (IF) | Developmental Studies Hybridoma Bank (DSHB) | GLA2A7 |
